# MeCP2-E1 isoform is a dynamically expressed, weakly DNA-bound protein with different protein and DNA interactions compared to MeCP2-E2

**DOI:** 10.1101/392092

**Authors:** Alexia Martínez de Paz, Leila Khajavi, Hélène Martin, Rafael Claveria-Gimeno, Susanne tom Dieck, Manjinder S. Cheema, Jose V. Sanchez-Mut, Malgorzata M. Moksa, Annaick Carles, Nick I. Brodie, Taimoor I. Sheikh, Melissa E. Freeman, Evgeniy V. Petrotchenko, Christoph H. Borchers, Erin M. Schuman, Matthias Zytnicki, Adrian Velazquez-Campoy, Olga Abian, Martin Hirst, Manel Esteller, John B. Vincent, Cécile E. Malnou, Juan Ausió

**Affiliations:** Department of Biochemistry and Microbiology, University of Victoria, Victoria, BC, V8W 3P6, Canada.; Unité de Mathématiques et Informatique Appliquées, Toulouse INRA, Auzeville BP 52627, 31326 Castanet-Tolosan cedex, France.; Centre de Physiopathologie de Toulouse Purpan, INSERM UMR 1043, CNRS UMR 5282, Université Toulouse III Paul Sabatier, Toulouse, France.; Institute of Biocomputation and Physics of Complex Systems (BIFI), Joint Units IQFR-CSIC-BIFI and GBsC-CSC-BIFI, Universidad de Zaragoza, 50018 Zaragoza, Spain.; Instituto Aragonés de Ciencias de la Salud (IACS), 50009 Zaragoza, Spain.; Aragon Institute for Health Research (IIS Aragon), 50009 Zaragoza, Spain.; Max-Planck-Institute for Brain Research, Synaptic Plasticity Department, Frankfurt/Main, Germany.; School of Life Sciences, École Polytechnique Fédérale de Lausanne, Brain Mind Institute, Lausanne, CH-1015, Switzerland.; Michael Smith Laboratories, University of British Columbia, Vancouver, BC, Canada V6T 1Z4.; Department of Microbiology and Immunology, University of British Columbia, Vancouver, BC, Canada V6T 1Z4.; University of Victoria-Genome British Columbia Proteomics Centre, Vancouver Island Technology Park, #3101-4464 Markham Street, Victoria, British Columbia V8Z7X8, Canada.; Molecular Neuropsychiatry & Development (MiND) Lab, Campbell Family Mental Health Research Institute, Centre for Addiction and Mental Health, Toronto, ON, M5T 1R8, Canada.; Institute of Medical Science, University of Toronto, Toronto, ON, M5S 1A8, Canada.; Department of Biochemistry and Microbiology, University of Victoria, Room 270d, Petch Building, 3800 Finnerty Road, Victoria, British Columbia V8P 5C2, Canada.; Gerald Bronfman Department of Oncology, Jewish General Hospital, Suite 720, 5100 de Maisonneuve Boulevard West, Montreal, Quebec H4A 3T2, Canada.; Proteomics Centre, Segal Cancer Centre, Lady Davis Institute, Jewish General Hospital, McGill University, 3755 Côte-Sainte-Catherine Road, Montreal, Quebec H3T 1E2, Canada.; Department of Biochemistry and Molecular and Cell Biology, Universidad de Zaragoza, 50009 Zaragoza, Spain.; Fundación ARAID, Government of Aragon, 50018 Zaragoza, Spain.; Biomedical Reseach Networking Centre for Liver and Digestive Diseases (CIBERehd), Madrid, Spain.; Michael Smith Genome Sciences Centre, BC Cancer Agency, Vancouver, BC, Canada V5Z 1L3.; Cancer Epigenetics and Biology Program (PEBC), Bellvitge Biomedical Research Institute (IDIBELL), Avinguda Gran Vía de L’Hospitalet 199-203. L’Hospitalet de Llobregat, Barcelona, Catalonia, Spain.; Physiological Sciences Department, School of Medicine and Health Sciences, University of Barcelona (UB), Catalonia, Spain.; Institució Catalana de Recerca I Estudis Avançats (ICREA), Barcelona, Catalonia, Spain.; Department of Psychiatry, University of Toronto, Toronto, ON, M5T 1R8, Canada.

## Abstract

MeCP2 – a chromatin-binding protein associated with Rett syndrome – has two main isoforms, MeCP2-E1 and MeCP2-E2, with 96% amino acid identity differing in a few N-terminal amino acid residues. Previous studies have shown brain region-specific expression of these isoforms which, in addition to their different cellular localization and differential expression during brain development, suggest they may also have non-overlapping molecular mechanisms. However, differential functions of MeCP2-E1 and E2 remain largely unexplored. Here, we show that the N-terminal domains (NTD) of MeCP2-E1 and E2 modulate the ability of the methyl binding domain (MBD) to interact with DNA as well as influencing the turnover rates, binding dynamics, response to nuclear depolarization, and circadian oscillations of the two isoforms. Our proteomics data indicate that both isoforms exhibit unique interacting protein partners. Moreover, genome-wide analysis using ChIP-seq provide evidence for a shared as well as a specific regulation of different sets of genes. Our findings provide insight into the functional complexity of MeCP2 by dissecting differential aspects of its two isoforms.

**Significance:** Whether the two E1 and E2 isoforms of MeCP2 have different structural and/or functional implications has been highly controversial and is not well known. Here we show that the relatively short N-terminal sequence variation between the two isoforms impinges them with an important DNA binding difference. Moreover, MeCP2-E1 and E2 exhibit a different cellular dynamic behavior and have some distinctive interacting partners. In addition, while sharing genome occupancy they specifically bind to several distinctive genes.

## Introduction

Methyl CpG binding protein 2 (MeCP2) was first identified through its ability to bind methylated DNA (1). Mutations in the *MECP2* gene were later associated with Rett syndrome (RTT; OMIM 312750), a severe neurological disorder that is among the most common causes of intellectual disability in girls (2). Affected females seem to develop normally during the first year but progression slows down until the age of 3-4. This is followed by a stagnation and subsequent regression that leads to the gradual loss of acquired communication and motor skills, eventually ending in a profound intellectual disability (2, 3).

*MeCP2* gene has four exons than can be alternatively spliced to produce two transcripts. The transcript skipping exon 2 has translation initiation in exon 1 and encodes MeCP2-E1. This isoform is slightly longer (498 amino acids in humans) and has 21 unique N-terminal amino acids. When exon 2 is included in the transcript, translation initiates in exon 2 to give rise to MeCP2-E2, a shorter variant (486 amino acids in humans) with 9 unique N-terminal amino acids (3, 4). The remaining sequence is identical for both isoforms, and encompasses the methyl binding domain (MBD), intervening domain (ID), transcriptional repression domain (TRD) and C-terminal domain (CTD) (5). MeCP2-E1 is likely the ancestral form of the protein, as orthologues are present across vertebrate evolution, whereas orthologous sequences of the exon 2 coding region have only been found in mammalian genomes (6).

Splicing variants often encode proteins with different functions, but in the case of MeCP2-E1 and E2 isoforms, this and the specific role played by the isoforms in the pathophysiology of Rett syndrome remain still controversial (7, 8). The presence of a polyalanine tract followed by a polyglycine tract in E1 NTD could be an indication of a potential functional difference (9). In this regard, polyalanine domains within various protein families are thought to have a convergent origin, suggesting that a specific function for these tracts might have been selected by evolutionary pressure (10). Potential evidence for the existence of non-overlapping functions of the E1 and E2 isoforms is supported by a difference in their relative abundance during development and in diverse regions of the brain (11, 12). MeCP2-E1 mRNA is the most abundant transcript in various regions of the brain (except for hypothalamus) (11), which, coupled to the reported inefficiency of translation of MeCP2-E2 relative to E1, may exacerbate their differential expression at the protein level (4). Moreover, Rett syndrome-causing mutations described so far involve solely the E1 isoform, and isoform specific mouse knockouts show Rett-related phenotypes for E1 knockout but not for E2, suggesting that E2 does not functionally compensate for the lack of E1 (13, 14). However, the high degree of structural similarity between MeCP2 isoforms point towards a high extent of functional overlapping, and some findings reinforce this idea. For instance, E2 expressed at levels comparable to those of E1 was reported to prevent key Rett-like phenotypes in mice models of Rett syndrome, indicating that part of the difference between isoforms could simply be related to the aforementioned disparity in temporospatial expression and protein levels (8).

Given the still controversial situation and the poorly understood nature of the differences between E1 and E2 isoforms, we decided to investigate this further. Our study comprehensively describes for the first time differences between MeCP2 isoforms, using various complementary biophysical, biochemical and genomic approaches. This work provides a detailed framework for the further understanding of the manifold functional aspects of MeCP2, thus shedding light onto the pathophysiology of Rett syndrome and other neurological disorders.

## Results

### MeCP2 E1 and E2 isoforms exhibit different cellular distribution and rate of expression during brain development

As it has been pointed out in the introduction, the issue regarding the different functionality of the MeCP2 isoforms has long remained controversial. However, there are many indirect hints to suggest otherwise. As Fig. 1 A clearly indicates, both isoforms exhibit quite a distinct neuronal localization. In hippocampal primary neuron culture the E1 antibody stains wide parts of the neuron and accumulation of immunoreactivity in the nucleus is not prevalent. Interestingly some MAP2-negative neurites – thus presumably axons – are labeled with the E1 antibody. This feature is also seen in the brain tissue section: although in the hippocampal CA3 region most E1 immunoreactivity is found as nuclear staining of pyramidal cell nuclei there is some immunoreactivity visible along the granule cell axonal projections in CA3 stratum lucidum. E2 staining in contrast is confined to few punctate structures in neuronal somata and nuclei in both hippocampal tissue and cultured cells, thus displaying a distribution pattern distinct from E1 immunoreactivity. Moreover, as seen in Fig. 1B, both E1 and E2 isoforms display a different pattern of expression during mouse brain development with E1 being present in a 15 fold excess to E2 at P15. Similar observations have been previously reported (12) and provide a framework for the structural and functional studies described in the following sections of this paper.

**Fig. 1.**
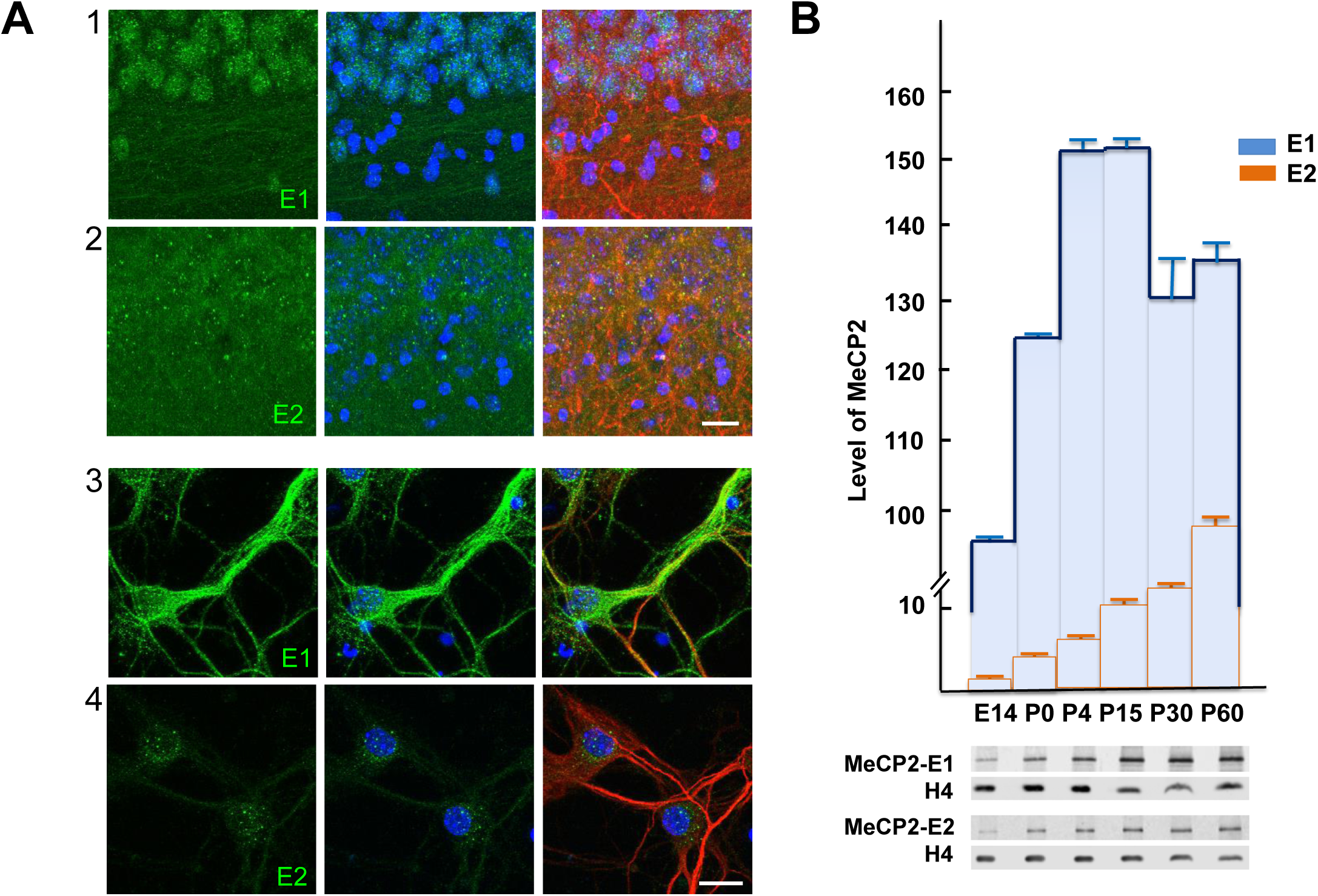
MeCP2 E1 and E2 isoforms exhibit distinct tissue and cellular distribution and different levels of expression during brain development. (A) *Staining pattern of MeCP2-E1 and E2 antibodies in mouse brain sections and hippocampal primary culture.* Confocal microscopic image close-ups of the hippocampal CA3 region from mouse brain sections (1, 2) and of cultured mouse hippocampal primary cells (3,4) showing antibody staining with E1 (1, 3) and E2 (2, 4) affinity-purified antisera (green). Nuclei are stained with DAPI (blue), neuronal dendrites and somata in with anti-MAP2 (red). Scale bars: 20 μm. (B) Western blot analysis of the changes in the level of expression of MeCP2-E1 and E2 isoforms during brain development.

### Biophysical characterization of MeCP2 isoforms N-terminal domains

The two MeCP2 isoforms differ only in their N-terminal domain (NTD) (Fig. 2 A), which has been previously described to lack any DNA specific binding structure but has the ability to stabilize the neighboring methyl binding domain (MBD) and its binding to methylated DNA (15, 16). A partial folding of this unstructured region might contribute to the interaction with double stranded DNA (dsDNA), thus having a differential impact on E1 and E2 binding properties. Therefore, we decided to compare the different biophysical properties of E1 and E2 NTDs. Constructs consisting of the E1 or E2 specific NTD followed only by the MBD were analyzed as previously described (16). Thermal unfolding studies of E1/NTD-MBD and E2/NTD-MBD were carried out by recording fluorescence emission as a function of temperature (Fig. 2 B). Estimation of apparent thermodynamic parameters for the structural stability was performed by fitting thermal denaturation curves according to the two-state unfolding model. Results indicate that E1 isoform shows a slightly lower mid transition temperature (*T*_*m*_) in all situations considered (Fig. 2 C; Supplementary Table 1), showing a slightly lower structural stability. This observation also includes the “closer to physiological” scenario (pH 7, 150 mM NaCl). Following the same trend, E1 isoform also shows a diminished unfolding enthalpy (Δ*H(T*_*m*_)), indicating a lower cooperativity in the thermal unfolding, suggesting that amino acid residues located at the NTD might be important for the stability of the folded regions located in the MBD. Furthermore, both isoforms are considerably stabilized upon addition of unmethylated or methylated dsDNA (a 45 bp fragment of dsDNA corresponding to BDNF promoter IV). The stabilizing effect is significantly larger in the presence of methylated DNA (Fig. 2 B and C). The nature of protein-DNA interactions was further assessed by determining their complete thermodynamic profile with isothermal titration calorimetry (ITC), considering a single binding site model (16) (Fig. 2 D and E). Results show that compared to E2, E1 exhibits 9-fold lower binding affinity (higher dissociation constant, *K*_*d*_) for methylated dsDNA and 5-fold lower binding affinity for unmethylated dsDNA, thus resulting in E1 isoform having a slightly lower discrimination capability for methylated/unmethylated dsDNA (Fig. 2 E). Strikingly, the main intermolecular DNA-binding driving forces for the two isoforms are of different nature, displaying opposed thermodynamic binding profiles: dsDNA interaction with E1 is enthalpically driven and with E2 is entropically driven; thus, while E1 interacts with favorable binding enthalpy (Δ*H*) and unfavorable binding entropy (-*T*Δ*S*), E2 interacts with negligible binding enthalpy and favorable binding entropy (Fig. 2 E). Therefore, while the interaction of E1 isoform with dsDNA is mainly driven by specific interactions between the protein and the dsDNA (*i.e.* hydrogen bonds and electrostatic interactions), the interaction of E2 isoform with dsDNA is mainly driven by unspecific interactions (*i.e*. hydrophobic desolvation and steric arrangements). In addition, E1 isoform exhibits a larger binding heat capacity (Δ*C*_*P*_) and the formation of its complex with dsDNA releases a larger number of protons (*n*_*H*_) Overall these observations indicate that the amino acid residues at the N-terminal regions of E1 and E2 NTDs have a strong influence not only on protein stability, but also on the interaction with the dsDNA: E1 is slightly less stable and exhibits lower affinity for dsDNA than E2 isoform. Fluorescence recovery after photobleaching (FRAP) data for the two isoforms supports this, with E1 having a more rapid recovery trajectory than E2, suggesting looser binding, although t-half and mobile fractions were not significantly different (Supplementary Fig 1). These properties could also be reflecting a differential ability of MeCP2 isoforms to interact with other molecules, namely proteins, nucleic acids or chromatin, as well as a different turnover rate, a different intracellular trafficking or differential susceptibility to undergo post-translational modifications.

**Fig. 2.**
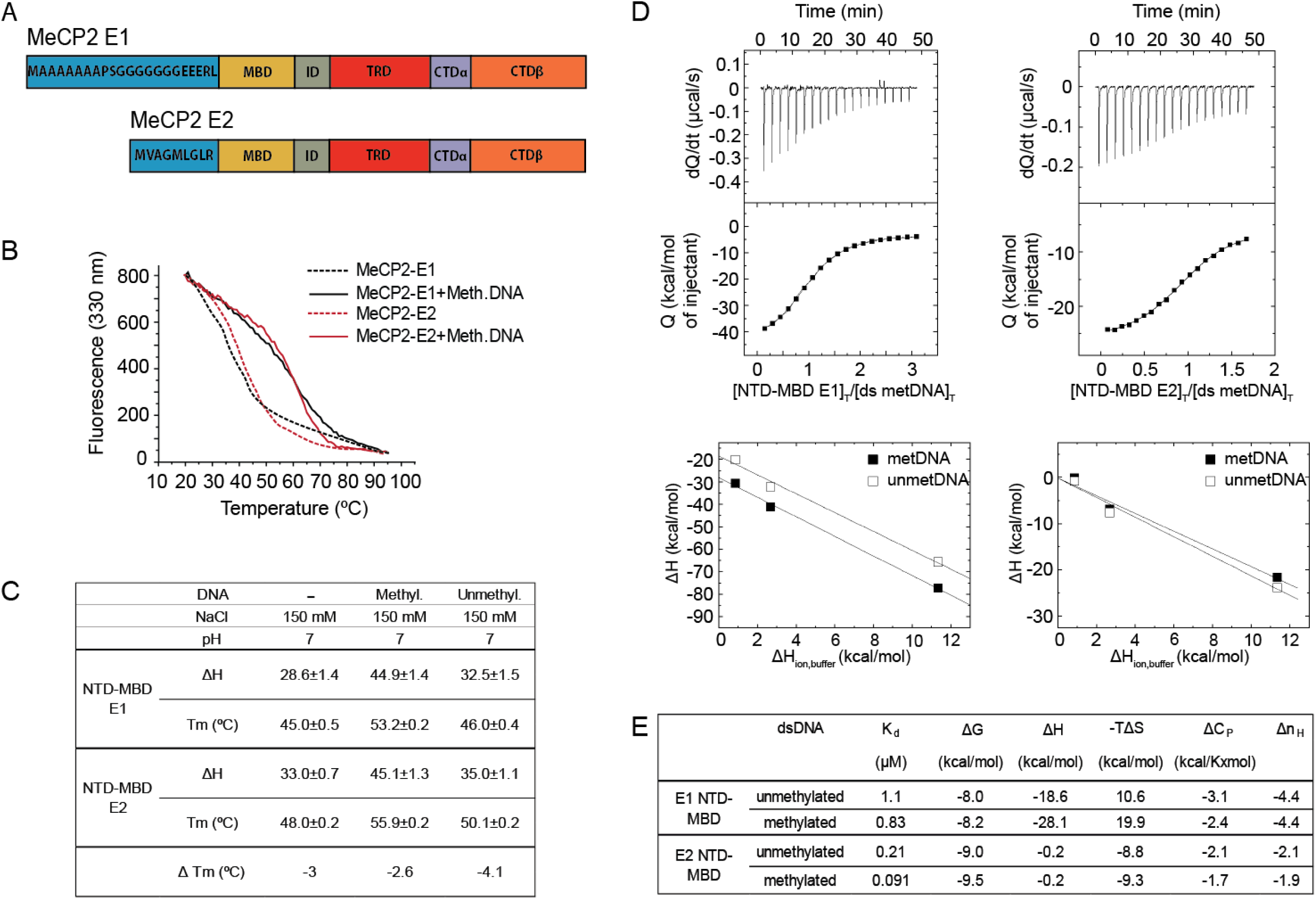
Biophysical characterization of the MeCP2-E1 and E2 NTD-MBD domains interaction with DNA. (A) Schematic representation of the MeCP2-E1 and E2 isoforms depicting the unique NTD amino acids sequences. (B) Fluorescence thermal denaturation curves for E1 and E2 NTD-MBD protein fragments in the presence of unmethylated and mCpG-dsDNA. All unfolding traces could be fitted considering a two-state unfolding model (not shown for clarity purposes). (C) Unfolding stability parameters obtained from thermal denaturations followed by intrinsic tryptophan fluorescence. (D) Calorimetric titrations of E1 and E2 NTD-MBD proteins interacting with dsDNA in Pipes (E1) and Tris (E2) 50 mM (pH 7) at 20 °C. The upper plots show the thermograms (thermal power as a function of time) and the binding isotherms (normalized heats as a function of the dsDNA/protein molar ratio). The use of buffers with different ionization enthalpy provided an estimation of the buffer-independent binding enthalpy and the net number of exchanged protons upon NTD-MBD-dsDNA complex formation by linear regression (lower graphs). (E) Buffer-independent dsDNA binding parameters (K_d_, dissociation constant; Δ*G*: Gibbs free energy of interaction; Δ*H*: enthalpy of interaction; -*T*Δ*S*: entropic contribution of interaction; Δ*C*_*P*_: heat capacity of interaction; *n*_*H*_: number of protons exchanged upon complex formation) obtained from calorimetric titrations at pH 7.

### Higher MeCP2-E1protein turnover in neuronal systems

MeCP2 is a member of the family of intrinsically disordered proteins (IDPs) (17). Such proteins are highly susceptible to proteolytic degradation, an important trait for proteins involved in dynamic cellular processes (18). IDPs can be protected from proteolysis by forming complexes with various molecules *in vivo* (18). The lower affinity of the E1 NTD-MBD region for DNA or its lower folding stability might reflect a higher presence in solution or the occurrence of a larger exposed surface to be targeted for proteasomal degradation (18). Therefore, we decided to compare the half-lives of the two MeCP2 isoforms in different neuronal systems. First, we performed transfections of undifferentiated SH-SY5Y neuroblastoma cells with E1- and E2-EGFP (enhanced green fluorescent protein) fusion proteins followed by cycloheximide (CHX) treatments-a compound that inhibits protein synthesis by blocking the peptidyl transferase activity of the 60S ribosome subunit. Western blot (WB) analysis of the CHX chase assays using an anti-EGFP antibody demonstrates that E1 has a faster turnover rate than E2. Approximately 50% of initial E1 is degraded 24 hours after CHX addition, while only 20% of E2 has been degraded by that time (Fig.3 A-1). Moreover, endogenous E1 and E2 levels were measured in similar way in SH-SY5Y induced to differentiation by a previously described procedure that uses a sequential treatment of retinoic acid and BDNF (19). Neuronal differentiation in these cells leads to the upregulation of MeCP2 expression (20) and to expression changes of differentiation markers (19) that were measured by WB and RT-qPCR respectively (Supplementary Fig. 2). Within this context we also observed a faster degradation of endogenous E1, around 30% of the initial E1 being degraded in 4 hours, while E2 level remains close to the initial protein amount at the same time (Fig. 3 A-2). To further confirm our findings, we followed a similar approach by performing CHX chase assays using DIV7 rat cortical neurons. Surprisingly, our E2 specific antibody was unable to recognize endogenous protein in these neurons. The E1 isoform showed a rapid turnover (40% of initial protein was degraded in 8 h after treatment initiation) (Fig. 3 A-3). Overall, these results point towards a significantly higher turnover rate for MeCP2-E1, suggesting a more dynamic role for this isoform.

**Fig. 3.**
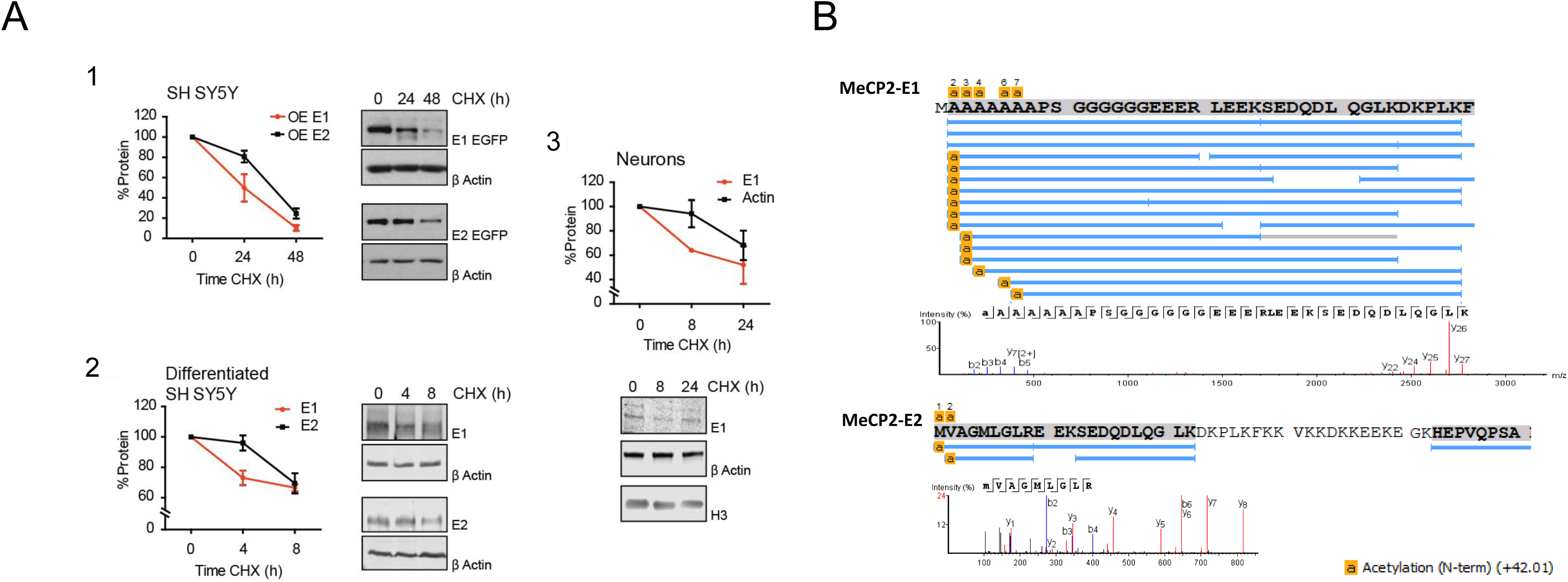
**(A)** Cycloheximide-chase assays of the E1 and E2 MeCP2 isoforms. Densitometries and representative western blots performed after cycloheximide treatments of (1) SH-SY5Y cells overexpressing (OE) E1 and E2 isoforms fused to GFP, (2) differentiated SH-SY5Y and (3) rat cultured cortical neurons with detection of endogenous E1 and E2 isoforms. Represented data are mean ± S.E.M. (n=3). MeCP2 levels were normalized using β actin and/or histone H3 as a control. **(B)** Mass spectrometry sequencing of the N-terminal end of the MeCP2 protein (*in vitro*). N-terminal peptide coverage alignment chart and high resolution mass spectra showing N-methionine excision (NME) and N-acetylation (NA) of the N-termini of MeCP2-E1 and MeCP2-E2. NA (+42 Da) of N-terminus amino acid is shown highlighted in yellow.

### Different N-terminal processing

To allow possible N-terminal *in vitro* modifications in a mammalian cell system, the NTDs of MeCP2-E1 and MeCP2-E2 were expressed in HEK293T cells and were subsequently purified using immunoprecipitation of recombinant fusion protein, followed by mass spectrometry and peptide analysis. Our MS analysis of PTM for MeCP2-E1 was reported previously (6), and showed no peptides with N-terminal methionine (NM), indicating complete NM excision (NME) at the first residue (P1) position. Acetylation of the initial alanine residue (P’1) after NME was observed (Fig. 3 B1). In addition, we observed some peptide reads with alanine 1, or alanine 1 and 2, or alanine 1 to 4, or 1 to 5 excised and acetylation of the subsequent alanine (Fig. 3 B). For MeCP2-E2, on the other hand, we found reads in which N-terminal methionine (P1 position) is retained and acetylated (Fig. 2B). For MeCP2-E2 we also found a few peptide reads with NME and acetylation of the penultimate valine (P’1) (Fig. 2B). All post-translational modifications (PTMs) reported received Ascores of 1000.

### Involvement of MeCP2-E1 in dynamic processes

MeCP2 is considered to modulate neuronal chromatin organization mainly through its ability to bind to methylated/hydroxymethylated DNA. Brain chromatin is a highly dynamic structure that undergoes remodelling changes that modify its accessibility upon neuronal activation (21) or other neuronal cues such as those involved in the circadian cycle (22). Likewise, DNA methylation appears to be more dynamic than previously anticipated and brain DNA methylation can be modified by activity (23, 24), and demonstrates 24-hour oscillations that resemble a circadian period (25). We previously described similar diurnal oscillation of MeCP2 protein levels and associated changes in chromatin accessibility in mice frontal cortex (26). This suggests that MeCP2 could be at the crossroads between DNA methylation oscillations and chromatin rearrangements resulting in gene expression changes upon different stimuli such as neuronal activation or circadian inputs. The higher abundance of MeCP2-E1 over E2 in neurons suggests that it could have a more prominent role in the regulation of these mechanisms. Hence, we decided to analyze the expression of MeCP2-E1 and E2 in two different settings. First, we took advantage of the system we reported previously displaying total MeCP2 24 h oscillations. We used frontal cortices of C57BL6 wild type mice euthanized at different time points during the day (26). MeCP2 function in frontal cortex appears to play an especially relevant role in Rett-syndrome as its levels within this tissue correlate with phenotypic severity in mice models of the disease (27, 28). Its ablation solely in forebrain neurons is associated with Rett-like behavioral impairments (29). Analysis of frontal cortices obtained at 12 a.m. and 12 p.m. show a noticeable 30% reduction of E1 protein level at 12 p.m., while E2 levels remain similar at these two times (Fig. 4 A). The second scenario involving MeCP2 dynamics was neuronal activation after KCl exposure. DIV7 rat cortical neuron activity was blocked by pre-treatment of the cells with tetrodotoxin (TTX), DL-2-amino-5-phosphonovalerate (APV) and 6-cyano-7-nitroquinoxaline-2, 3-dione (CNQX). Neuronal depolarization was subsequently achieved by 30 minute exposure to 55 mM KCl and the protein levels were measured at different time points after depolarization. The inability of the MeCP2-E2 antibody to detect this isoform in rat neurons prompted us to firstly determine the endogenous levels of total MeCP2 after treatment. Due to the high E1 abundance compared to E2, this mostly corresponds to the E1 isoform. Next we assessed the two isoforms’ dynamics by transfecting cultured rat neurons with flag-tagged E1 and E2 constructs. The results of these two approaches show a fast increase of total MeCP2 levels immediately after KCl treatment followed by a decrease to basal levels at around 4 hours after treatment [Fig. 4 B; all points normalized to NT (non-treated) samples (value = 1)]. Interestingly, we observe a completely different pattern between MeCP2 isoforms. As expected, E1 shows a trend similar to that of total MeCP2, rapid upregulation upon depolarization that is maintained during 3-4 hours, and then protein levels decrease to reach, in this case, approximately 50% of the initial E1 levels (Fig. 4 C). By contrast, E2 shows a stable pattern, exhibiting levels which are similar to those of non-treated cells throughout the whole duration of the experiment (Fig. 4 C). Hence, our data confirm the existence of different dynamics of MeCP2 isoforms in the two different settings studied, and are consistent with a different role of the two MeCP2 isoforms within the neuronal context.

**Fig. 4.**
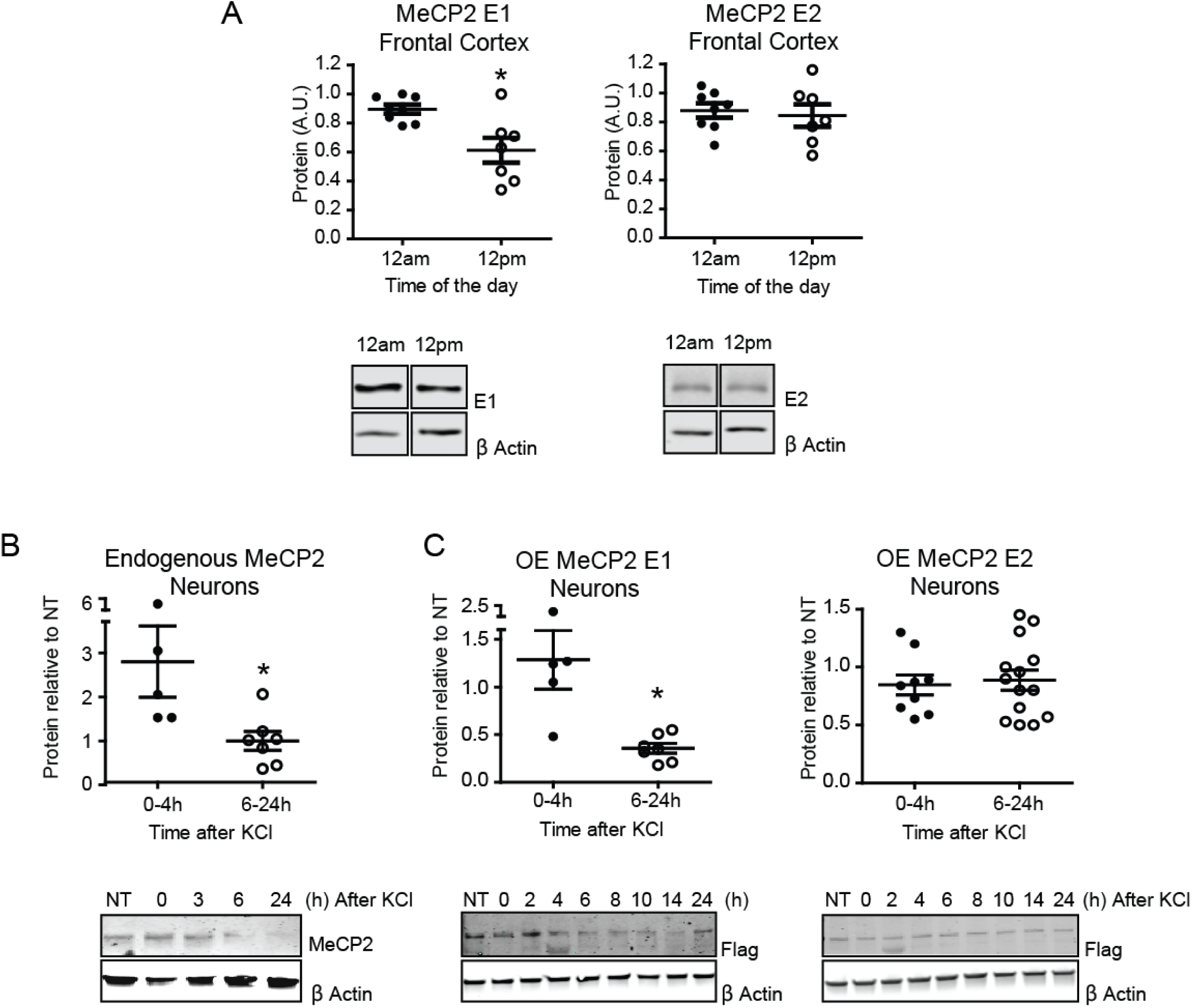
MeCP2 isoforms dynamics during the day and upon neuronal depolarization. Densitometric analysis and representative western blots data showing: (A) endogenous E1 and E2 levels in frontal cortices of mice euthanized at 12 a.m. and 12 p.m., (B) total endogenous MeCP2 of cultured cortical neurons upon KCl depolarization and (C) E1 and E2 levels of transfected cultured cortical neurons overexpressing (OE) Flag-MeCP2-E1 or E2 upon KCl depolarization. NT represent non-treated cells and all values are normalized to this point (value = 1). Represented data are mean ± S.E.M (n=3-4). * *P* < 0.05 Two tailed Mann Whitney test. MeCP2 levels were normalized using β actin and/or histone H3 as a control.

### Genome wide distribution of MeCP2-E1 and MeCP2-E2 isoforms

The differences described between E1 and E2 in terms of their affinity for methylated DNA and daily dynamics might have an influence on and/or reflect a differential genomic distribution. Therefore, we decided to investigate this possibility by performing chromatin immunoprecipitation sequencing (ChIP-seq) analysis of E1 and E2 in frontal cortices of mice euthanized at 12 a.m. and 12 p.m. As it has been previously noted for total MeCP2 (30, 31), both isoforms exhibited a very broad chromatin binding pattern (Fig. 5 A). Consequently, the reads obtained from the sequencing of two independent biological replicates for each time-point were merged in order to enhance any slight differences between the IPs and the input samples. IP reads were then normalized to their respective inputs (log2 ratio). We used spatial clustering for the identification of ChIP enriched regions (SICER) (32). This analysis indicated that the overall distribution along different genomic regions was similar, with half of the peaks called for both E1 and E2 overlapping with intergenic regions (FDR ≤ 0.001, SICER algorithm: window 600 bp; gap 200 bp; Fig. 5 B). In agreement with the daily changes observed in E1 protein levels at 12 p.m. (Fig. 4 A), the number of E1 enriched regions at this time also decreased (4242 at 12 a.m. *vs.* 3052 at 12 p.m.), while a more modest difference was observed for E2 (2371 islands at 12 a.m. *vs.* 2108 at 12 p.m.) (Supplementary Fig 3A). Occupancy differences over time were more pronounced if we took into consideration the levels of E1 and E2: E1 levels decreased in 1490 regions and increased in 575 while E2 decreased in 635 regions and increased in 434; (Supplementary Fig. 3 B), with the biggest variations found at intergenic regions for both isoforms (Supplementary Fig. 3 C). Despite the similar isoform’s general distribution, we were able to identify different significantly enriched binding motifs for both isoforms. Using the RSAT tool we detected distinctive motifs: ATACAC (p-value: 4.4e-11) and CCACAG (p-value: 3.9e-12) for E1 and CAAAAC (p-value: 3.4e-3) and CAAAAG (p-value: 2.4e-3) for E2, indicating a differential binding site preference (Fig. 5 C).

**Fig. 5.**
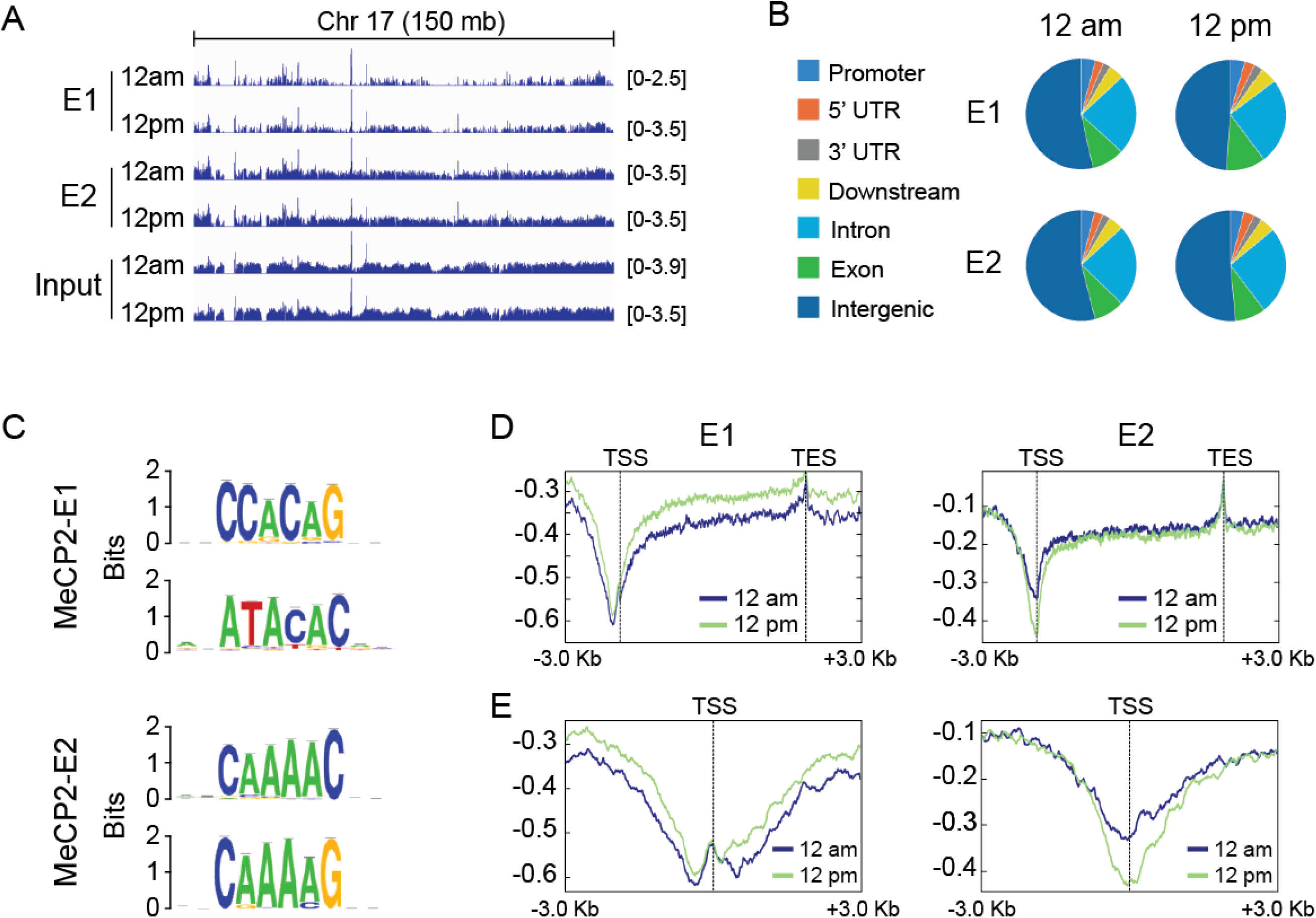
Genome wide distribution and dynamics of MeCP2 isoforms. (A) Integrative genomics viewer (IGV) (80, 81) profiles of E1 and E2 ChIP-seq (first four rows E1 12 a.m. and 12 p.m. followed by E2 12 a.m. and 12 p.m.) and Input DNA (last rows input 12 a.m. and 12 p.m.) based on sequence reads. Data represent a 150 Mb region of chromosome 17. Numbers in the right of each track represent track heights (B) Pie charts depicting the distributions of regions enriched in the MeCP2 isoforms [as detected by Spatial clustering for the Identification of ChIP Enriched Regions (SICER); window 200 bp and gap 600 bp] across 7 defined genomic features: promoters, 5’ UTR, 3’ UTR, proximal downstream of genes, introns, exons and intergenic regions. (C) DNA-binding preferences E1 (upper graphs) and E2 (lower graphs) shown as motif logos based on aligned, over-represented patterns found in the binding sites. The overall height of each stack indicates the sequence conservation at that position (measured in bits), whereas the height of symbols within the stack reflects the relative frequency of the corresponding amino or nucleic acid at that position. (D) Summary profiles across 3 Kb upstream the TSS and 3 Kb downstream the TES of genic regions occupied by E1 at 12 a.m. and 12 p.m. (left graph) and E2 at 12 a.m. and 12 p.m. (right graph). (E) Detail of the previous representations focusing in a 6 Kb region surrounding the TSS of E1 and E2 bound genes (left and right panels respectively).

As MeCP2 has been described to be a transcriptional regulator, we decided to analyze the distribution of MeCP2 isoforms around transcribed regions. The summary binding profiles of E1 and E2 to regions spanning 3 kb upstream of transcription start sites (TSS) to 3 kb downstream transcription end sites (TES) demonstrate a consistently similar binding pattern for both isoforms, with a marked depletion at TSS and a peak at TES (Fig. 5 D). Moreover, there is a slight decrease in E1 at 12 p.m. in its binding to gene bodies while E2 exhibits a slight decrease around the TSS. Interestingly, a closer inspection around TSS regions revealed that the E2 isoform displayed a marked depletion precisely at the TSS. In contrast, the E1 isoform is depleted at both sides of the TSS corresponding to the +1 and −1 nucleosome regions, with a slight increase on the TSS. These results suggest a differentiated role of the two isoforms in shaping the chromatin structure around the TSS.

We then clustered the genes based on their MeCP2 occupancy using deepTools (33) Heatmap clusters and profiles for the log2 ratio plots failed to reveal any differential binding of the isoforms to specific gene clusters (data not shown), however we detected daily differences for each isoform occupancy throughout gene bodies (Fig. 6 A). For instance, E1 cluster 1 showed a flat profile at 12 a.m. and an increased binding at 12 p.m. (Fig. 6 B). In the case of E2, cluster 4 exhibited an increased binding at 12 p.m. compared to 12 a.m., while cluster 5 displayed lower binding at 12 p.m. (Fig. 6 C). Functional pathways associated with genes present in each cluster were analyzed using the Kyoto encyclopedia of genes and genomes (KEGG) (Fig. 6 D left graphs). All three clusters were enriched in genes related to sensory transduction like olfaction or taste, and with histone proteins (E1 was associated with genes encoding H2A family members (p-value: 1.16 e^−06^) and E2 mainly with members of the histone cluster 1 (cluster 4 p-value: 1.65 e^−13^ and cluster 5 p-value: 2.23 e^−27^). MeCP2 isoform-specific enrichments were related to neuroactive ligand-receptor interaction in E1 and ribosomal proteins in E2. Interestingly, cluster 5 contain several genes associated with the neurodegenerative diseases Huntington (p-value: 4.17 e^−08^), Parkinson (p-value: 9.87 e^−06^) and Alzheimer (p-value: 9.15 e^−06^). ChIP-qPCR validations of randomly selected genes of each cluster confirmed the general trends observed in our ChIP-seq-analysis, despite the very slight variations of the isoforms occupancies during the day (Fig. 6 D right graphs). Overall, our results suggest that beyond the common functions in which both isoforms are involved, they regulate different sets of genes and display distinct dynamics on their genomic occupancy, reinforcing the existence of non-overlapping roles.

**Fig. 6.**
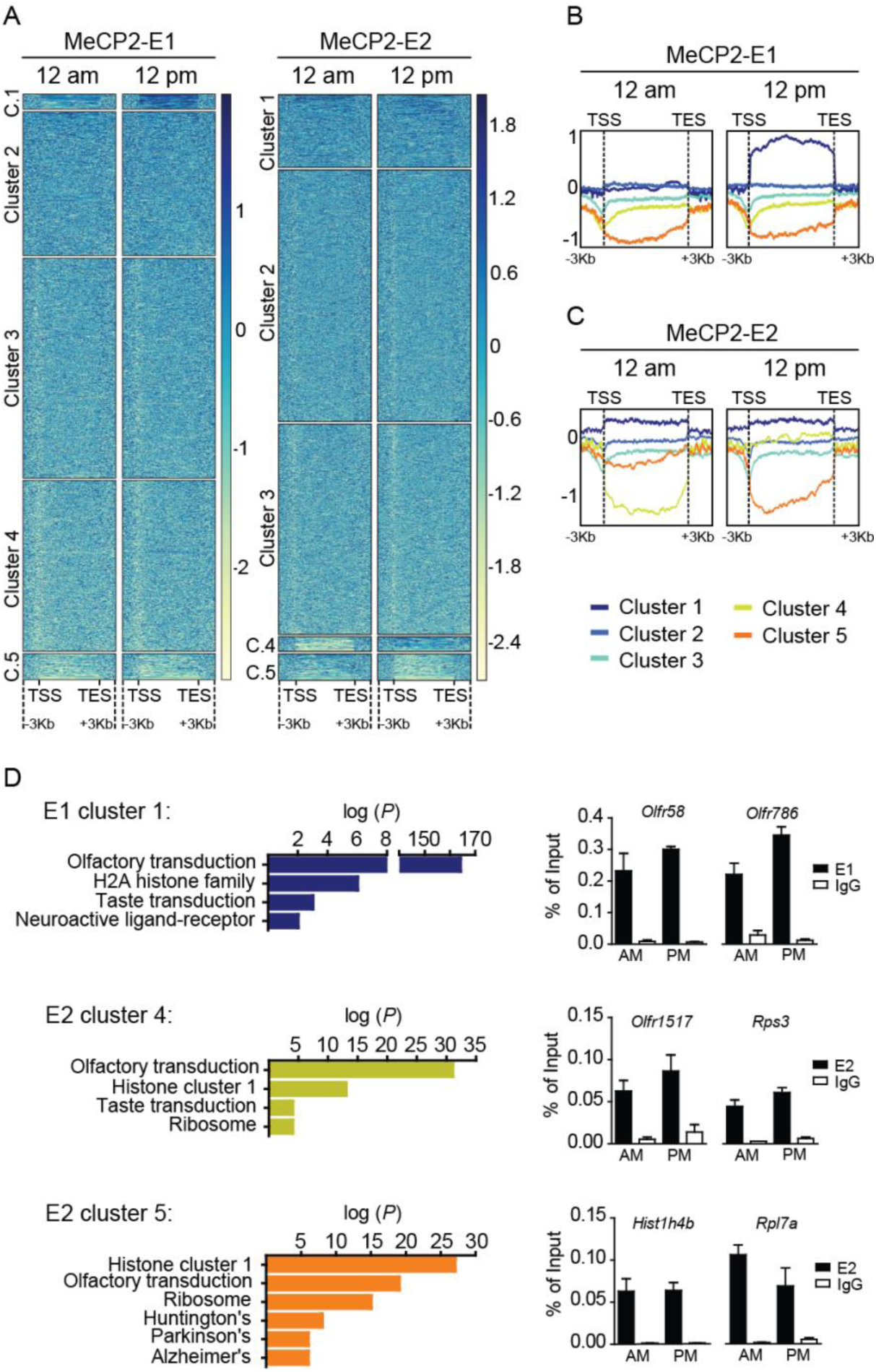
MeCP2-E1 and E2 isoforms display diurnal dynamic genomic binding. (A) Heatmaps representing the log2 ratios obtained for E1 and E2 ChIP experiments, where each row is a gene and each column is divided into five clusters using the k-means algorithm. Protein occupancy is represented by color intensity, where the darker the color, the higher the protein enrichment. (B) Comparison of E1 enrichment at 12 a.m. *vs*. 12 p.m. showing occupancy differences in different clusters of interest (1: dark blue; 2 blue; 3: cyan; 4: yellow; 5: orange). (C) Heatmap depicting the E2 12 a.m. vs. 12 p.m. shows a dynamic binding in clusters 4 and 5 (yellow and orange respectively). (D) Left graphs: top-enriched functional pathways (-log10 (*P*)) of genes included in dynamic E1 and E2 gene clusters [Kyoto encyclopedia of genes and genomes (KEGG)]. Right graphs: ChIP-qPCR results validating MeCP2-E1 and E2 variations in occupancy of genes included in each of the gene clusters obtained.

### MeCP2-E1 and E2 protein partners

IDP proteins are characterized by their inability to acquire a stable secondary structure when free in solution. This confers the structural flexibility that enables them to serve as scaffolds for the recruitment of partners and thus function as interacting hubs (34). Interestingly, IDPs, including MeCP2, usually acquire ordered structures upon binding to their interacting partners, allowing the exposure of molecular recognition features (MoRFs) to further make contacts with other molecules (35, 36). Thus, the possibility exists that the aforementioned E1 and E2 differences in unfolding temperature and affinity for DNA could expose differential interacting surfaces. These attributes together with their previously discussed expression patterns (3, 4, 37) raise the possibility that E1 and E2 might be involved in non-overlapping molecular functions that perhaps could be defined through the identification of their protein interactors. Therefore we decided to perform a comprehensive proteomic analysis to look for MeCP2-E1 and E2 protein partners. We performed this analysis on mice whole brain nuclei (Fig. 7 A), and chromatin was extensively digested with micrococcal nuclease (MNase) to release as much MeCP2 as possible, including that which could be embedded in tightly condensed chromatin regions. Endogenous E1 and E2 were subsequently immunoprecipitated from lysates of the MNase digested nuclei with antibodies specific for each of the two MeCP2 isoforms (Supplementary Fig. 4 WB and IP E1 E2). Normal rabbit IgG and blocking of E1 and E2 antibodies with blocking peptides were used as negative controls. Co-immunoprecipitated proteins were separated by SDS-PAGE and different gel fractions sectioned for protein identification by mass spectrometric analysis (Fig. 7A).

**Fig. 7.**
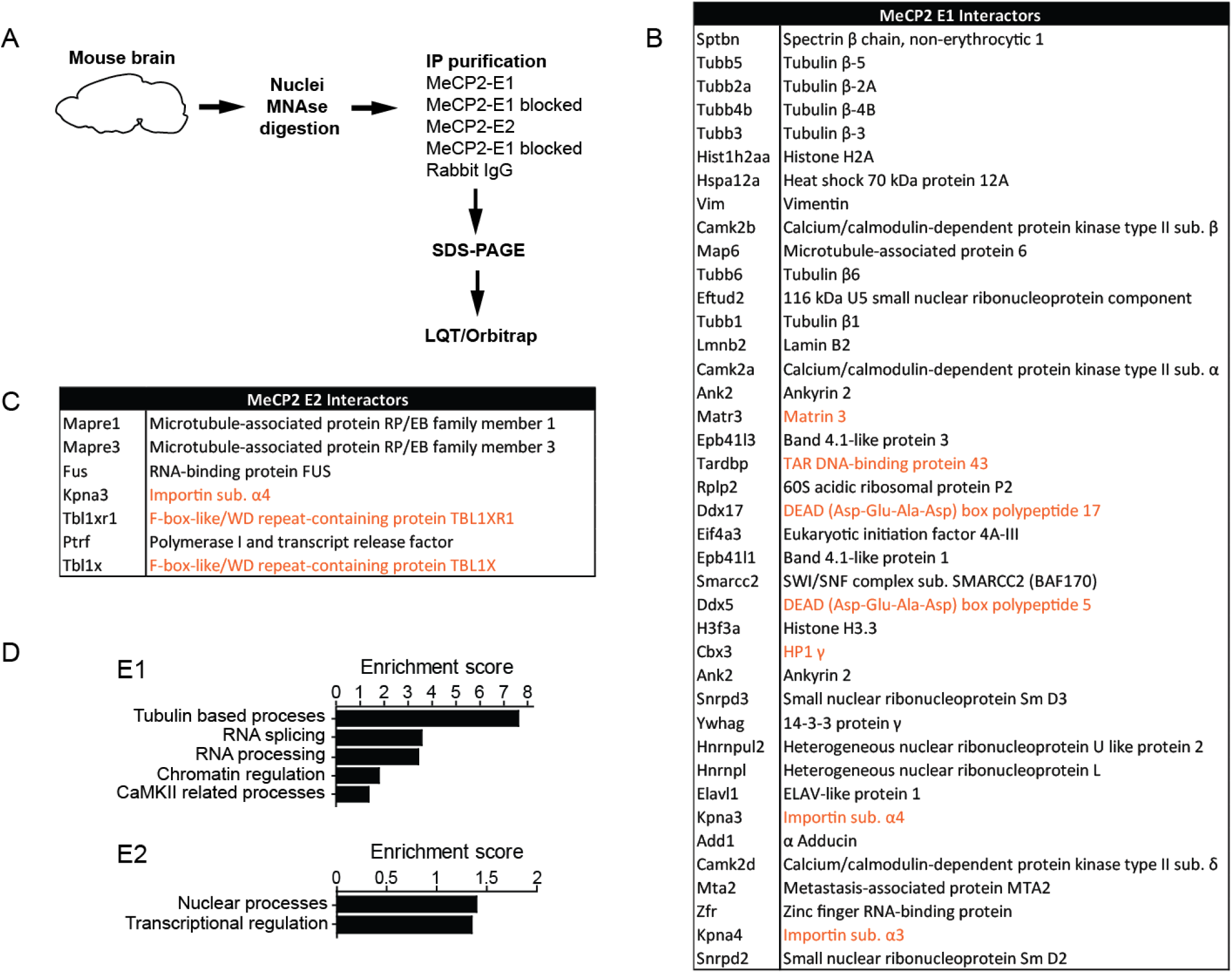
MeCP2-E1 and E2 interacting proteins. (A) Schematic workflow of the proteomic analysis. (B and C) Proteins partners identified for each isoform. (D) Selected pathways enriched in E1 and E2 interacting proteins as identified by functional clustering (DAVID Gene Ontology Bioinformatics Resources). Dotted line, *P*<0.05 from EASE score, a modified Fisher’s exact *p*-value.

We chose proteins identified by at least 2 significantly matching peptides which were absent from the negative controls. According to our expectations this filter rendered a great number of potential interacting proteins for the E1 isoform, 40, and 7 for E2 (Fig. 7B and 7C). As a good validation for our approach we detected several previously described MeCP2 interactors (Fig. 7B and 7C, interactors highlighted in orange (38-44)). Functional clustering of co-eluted proteins (DAVID (45)) uncovered functional enrichments, especially for E1 (Fig. 7 D). E1 co-eluted proteins are highly enriched for β-Tubulins, the building blocks of microtubules, and microtubule-associated proteins such Adducin 1 (Add1) or microtubule associated protein 6 (Map6). Importantly, microtubule assembly initiates from the centrosome, organelle associated to MeCP2 function in microtubule stability and mitotic spindle organization (46-48). Proteins related to mRNA splicing and mRNA processing were also highly represented among E1 partners (for example 116 kDa U5 small nuclear ribonucleoprotein component [Eftud2], Heterogeneous nuclear ribonucleoproteins L [Hnrnpl] or DEAD (Asp-Glu-Ala-Asp) box polypeptides 5 and 17 [Ddx5 and Ddx17]). MeCP2 functions on RNA splicing or mRNA processing have been previously described (43, 49, 50) but still lack deep investigation. As we expected, functions related to chromatin regulation are also enriched among MeCP2-E1 partners, as we found the nucleosome-core histone H2A and the variant H3.3, the chromatin regulators Brg1 associated factor 170 (BAF170), member of the switch/sucrose non fermenting (SWI/SNF) complex, and MTA2, subunit of the nucleosome remodeling deacetylase (NuRD) complex (51). Functional network analysis (STRINGv10 (52)) revealed a higher than expected number of connections between all E1 and E2 interactors (Supplementary Fig. 5; p-value<0.001) and suggests the participation of E2 in processes similar to those involving E1, but through the interaction with a different set of protein partners. In this regard, among E2 interactors we found the microtubule-associated protein RP/EB family members 1 and 3 (Mapre1 and Mapre3), important for microtubule organization (53). The E2 interactor fused in sarcoma (FUS) is involved in mRNA processing, with *Mecp2* being one of its known target genes (54). In the chromatin regulation function we found that E2 specifically interacts with two recently described MeCP2 protein partners: Transducin-β-like 1 (Tbl1) and Tbl1-related 1 (Tbl1r1), components of the nuclear receptor co-repressor (N-CoR) complex (38, 39). Interestingly, E2 also interacts with the polymerase I transcription and release factor (Ptrf), protein involved in ribosomal DNA (rDNA) transcription through the formation of the rDNA loops (55).

E1 co-eluted proteins include spectrin β1, lamin B2, the band 4.1 proteins B and N and matrin 3 (the latter was previously reported to interact with E1 in neuronal nuclei (41)), components of the nuclear matrix (56), classically defined as a fibrogranular structure which consists of nucleoskeleton/nuclear lamina networks and associated proteins (57, 58). Of note, one of the best characterized components of the nuclear matrix is the attached region binding protein (ARBP), a chicken MeCP2 orthologue (59) that binds methylated DNA within matrix attachment region (MAR) elements (58, 60). Overall, the lack of shared protein partners by the MeCP2-E1 and E2 isoforms suggest their involvement in similar general mechanisms like RNA processing, chromatin control of transcription or microtubules regulation, but performing non-redundant functions through the interaction with different partners.

## Discussion

The existence of mutations affecting only the MeCP2-E1 isoform in Rett patients [*e.g.* p.Ala2Val; (14, 61)] suggest that endogenous E2 expression cannot compensate for the lack of functional E1. An important question thus arises as to whether this is simply related to the lower levels of E2 found in neurons (37) or it is due to the existence of E1 specific functions that cannot be provided by the E2 isoform. The different cellular distribution of the two isoforms and their distribution during brain development (Fig.1) also suggest a different functionality.

The NTD is the only structural feature that differs between the two MeCP2 isoforms, and currently there is a lack of information regarding any potential functional difference between E1 and E2 NTDs. The NTD of these isoforms has generically been described as a highly disordered region able to acquire secondary structure, as demonstrated by the coil-to-helix transitions exhibited in the presence of hydrogen bond stabilizers (36). Such conformational transitions contribute to enhancing the MBD affinity for DNA (15). To the best of our knowledge, the biophysical characterization of the interaction of NTD-MBD fragments of the two MeCP2 isoforms with DNA is the first of its kind (Fig. 1). Most of MeCP2 functions rely on its ability to bind nucleic acids, and in this regard these results uncover a fundamental 9-fold difference in affinity for DNA of E2 over E1 isoform. This could be one of the basic structural features responsible for shaping the functional discrepancies between the isoforms.

The main differences observed here between the MeCP2 isoforms can be summarized as follows: E1 (the major isoform of MeCP2 in neurons) shows a lower DNA binding affinity and a lower structural stability (Fig.1). E1 also exhibits a higher basal degradation rate in various neuronal settings (Fig. 3), and enhanced dynamic fluctuations of protein levels *via* diurnal rhythm oscillations and neuronal depolarization (Fig. 4). Within a neuronal context these attributes are especially interesting given the peculiar chromatin relationship between MeCP2 and the linker histone H1 in this tissue. Under normal conditions, neurons express MeCP2 to near histone-octamer levels and contain half the amount of H1. By contrast, when MeCP2 is not expressed, H1 returns to the levels which are observed in other somatic tissues (62). Therefore, it is likely that MeCP2-E1 could function within this setting as a DNA-methylation dependent highly dynamic linker histone, needed to allow for rapid chromatin structural changes in response to external stimuli. This is particularly important in neurons, given their versatile ability to readily modify gene expression as a result of their unique methylome (24, 63).

In this regard, signal transduction from the cell surface to the genome often relies on the cytoskeleton-nucleoskeleton-chromatin interconnection (56). Importantly, we have identified MeCP2-E1 and E2 protein partners associated to every part of this system: the cytoskeleton (i.e. tubulins, Map6, Mapre1 and 3), nuclear envelope/matrix associated proteins (Lamin B2, Band 4.1 proteins, Spectrin or Matrin 3) and chromatin (histone proteins, Hp1γ, Mta2 or Baf170). Our findings open up the possibility of MeCP2 functioning as an important player in signal transduction. In particular, E1 could play a prominent role in the neuron-specific nucleoskeleton-chromatin connection, due to its remarkable abundance and the dynamism observed upon the application of external stimuli such as neuronal depolarization.

Because of its higher abundance, E1 appears to exhibit a more dynamic behavior, however, E2 also exhibits an oscillating genomic binding nature. Surprisingly, most of the daily MeCP2 isoform binding differences observed overlap with genes encoding for sensorial receptors, such as olfactory (ORs) and taste (TASRs) receptors (Fig. 6 D). This seemingly counterintuitive result is very interesting as preliminary observations have shown expression of ORs and TASRs in brain regions not related to the direct detection of odors and flavors (64). The study of the so called “ectopic ORs” (outside olfactory epithelium) is in its infancy, but apparently they act as chemoreceptors which are important to maintain cellular homeostasis, and some of them are able to activate complex cellular responses mediated by neurotransmitters or hormones (64). The expression of olfactory receptors has been described to be upregulated in MeCP2 KO mice and downregulated in mice overexpressing the protein. These data support a potential role for MeCP2 in transcriptional regulation of these genes in different regions of the brain structures such as cerebellum, amygdala and hypothalamus (30). More importantly, the expression of these receptors in brain is altered in neurodegenerative diseases such as Parkinson’s, Alzheimer’s and in prefrontal cortex in schizophrenia (64), disorders in which MeCP2 expression has been also observed to be dysregulated (17). Our ChIP-seq results reinforce this MeCP2 function and add a dynamic component to it.

Quite unexpectedly, all gene clusters displaying dynamic diurnal binding of the two MeCP2 isoforms were also enriched in genes encoding replication-dependent (RD) histones. Expression of RD histones has been recently detected in terminally differentiated cells and tissues, including neurons and brain (65, 66). Some of these genes encode histone isotypes with a certain sequence divergence (65), and could possibly affect the histone interactions within the nucleosome. Further analyses will be required to assess if MeCP2 has any regulatory effect on the expression of such genes, but it is tempting to speculate a possible role in the generation of variant nucleosomes present in adult brain (65).

An additional distinctive function for MeCP2-E2 which could be inferred from our results has to do with ribosomal gene expression regulation. Our co-immuno-precipitation experiment demonstrated the interaction of E2 with Ptrf (Fig. 2). Transcription of ribosomal genes has been reported to be dependent on the DNA methylation status (67) and occurs through the formation of nucleotide loops linking initiation and termination gene regions, process in which Ptrf participation is essential (55). Our ChIP-seq results provide evidence for a dynamic MeCP2-E2 genomic binding to ribosomal genes (Fig. 6). In support to our results, MeCP2 has been previously linked to nucleolar changes during neuronal maturation (68).

Another observation made through our ChIP-seq analysis is that the two isoforms possess significant differences in DNA-binding site preference; E1 targets are significantly enriched for the DNA motifs ATACAC and CCACAG, whereas E2 targets are enriched for CAAAAC and CAAAAG.

Overall, the present work provides support to the notion that Rett syndrome arises from the simultaneous impairment of different cellular functions involving both MeCP2 isoforms. For instance, mutations of E1 may have a larger impact in neuronal chromatin structure and stimuli-dependent gene expression dynamics. We have previously described in *MeCP2* KO mice a decrease in in the circadian gene oscillations of the brain derived neurotrophic factor (*Bdnf*) and Somatostatin (*Sst*) genes (26). By contrast, similar mutations in E2 could be involved in the deregulation of ribosomal expression or microtubule control.

The seemingly contradictory literature available to date, regarding the degree of functional overlapping of MeCP2 isoforms is likely the result of the lack of studies carried out on the endogenous native proteins. Overall our results provide strong support for the existence of both different and overlapping functions between the two. Importantly, our study uncovers some functional aspects of MeCP2 that were previously unknown. It opens the door to further investigation that will be helpful to understand the role of this complex protein in the healthy state and the consequences of its deregulation in Rett syndrome and other neurological disorders.

## Materials and Methods

### Animals

C57BL/6 mice were maintained under standard animal house conditions (12 hour dark-light cycles on *ad libitum* food and water intake). The experimental procedures were in accordance with all legislation defined by the European Union and approved by the local ethics committee (UB-IDIBELL). For preparation of rat neuron primary culture used in Figures 3 and 4, animal manipulations were performed in agreement with the European Union Council Directive 86/609/EEC and following the French national chart for ethics of animal experiments (articles R 214-87 to 90 of the “Code rural”). Preparation of primary neuron received approval from the local ethic committee (permit number: 04-U1043-DG-06).

### Immunostaining

The distribution pattern of the affinity-purified antibodies was tested in mouse hippocampal primary cultures and in mouse brain sections. Primary cultures were prepared from P0/P1 mouse pups as described earlier (69), cultured for 7 days in glass bottom dishes, fixed for 20 min with 4% para-formaldehyde/4% sucrose in PBS, permeabilized 15 min with 0.5% Triton in BB (PBS with 4% goat serum), blocked in BB for 1h and incubated with primary antibodies in BB for 1 h. After washing, secondary antibodies in BB were applied for 30 min in the presence of 1 μg/μl DAPI to stain nuclei. Cells were washed and imaged in PBS. For mouse brain slices 50 µm cryostat sections were cut from 4% PFA-fixed, adult male mouse brain which was dissected after deep anesthesia and decapitation, fixed and cryoprotected in increasing concentrations of sucrose before freezing in OCT and sectioning. Sections were postfixed for 15 min and stained free floating by permeabilization and blocking in 0.5% Triton in BB for 2 hrs at RT and incubation with primary antibodies in 0.5% Triton in BB overnight at 4°C. After washing secondary antibodies were applied for 2 hrs in 0.5% Triton in BB at RT together with 1ug/ul DAPI to stain nuclei. Sections were mounted on slides with Aquapolymount (Polysciences). Antibodies were used in the following dilutions: E1affi 1:200 (cells), 1:250 (sections), E2 affi 1:100 (cells), 1:250 (sections), MAP2 (gp) 1:1000 (Synaptic Systems), secondary antibodies produced in goat coupled to Alexa488 and Alexa594 dyes were used 1:1000 (Life Technologies/Thermo Fisher Scientific). Imaging was performed on a Zeiss LSM780 confocal microscope (Zeiss, Oberkochen) with a Plan-Apochromat 40x/1.4 Oil DIC objective (cultured cells) and a Plan-Apochromat 20x/0.8 (WD: 0.55 mm objective (tissue section) as Z stacks. Z-stacks were converted in the ZEN software (Zeiss) to maximum intensity projections. Images were adjusted for brightness and contrast and close-ups were cropped from the entire image in ImageJ to visualize details of the CA3 region and the cultured cells.

### Analysis of MeCP2-E1 and MeCP2-E2 variation during brain development

Three female mouse brains were pooled and homogenised for each development time period to prepare the samples for western blot analysis. Samples were prepared as per (70) and stored at −80°C until use. For each time point, western blots were repeated at least 6 times to get the average densitometry values. Western blots were carried using in house antibodies for MeCP2-E1 and E2 (see supplementary materials and methods and supplementary Fig.3; see also next section). In contrast to similar data published in (12) we used a nuclear protein (histone H4) as a normalizer also because actin has been shown not to be an appropriate candidate for studies related to neuronal differentiation (71). To quantify the ratio of MeCP2-E1/MeCP2-E2 we used the nuclear extracts of HEK293 cells expressing the 3 x Flag versions of these isoforms (See Supplementary Fig. 4). Stocks of these extracts containing an identical amount of (MeCP2) protein as determined by Western and Li-Cor quantification (see below) using a C-terminal MeCP2 antibody were prepared. These stocks were used to prepare serial dilutions to produce linear plots against which the amounts of native MeCP2-E1 and E2 were measured and their ratio determined.

### Western blot

Frozen frontal cortices and cell pellets were homogenized on Laemmli buffer [2% SDS, 10% glycerol, 0.002% bromophenol blue and 62.5 mMTris-HCl (pH 6.8)]. Protein content was measured by the Lowry method (Bio-Rad). Immediately after, 3% B-mercaptoethanol was added, proteins were separated by sodium dodecyl sulfate polyacrylamide gel electrophoresis (SDS-PAGE) and transferred onto nitrocellulose membranes (GE Healthcare). Nonspecific binding was blocked by incubation in 5% skimmed milk in phosphate buffered-saline (pH 7.2) with 0.1% Tween 20. Membranes were incubated with specific antibodies, either overnight at 4°C or for 1 hour at room temperature. Antibodies and dilutions used were as follows: MeCP2 1:5000, β-actin 1:20000, Histone H3 1:20000 (Abcam), Flag 1:5000 (Sigma-Aldrich), EGFP 1:1000 (Thermo Fisher Scientific), in house rabbit polyclonal antibodies MeCP2 E1 1:1000 and E2 1:500. Fluorescent secondary antibodies (Li-Cor) 1:10000. Densitometries were performed with Li-Cor Image Studio Light software.

### Protein expression

Plasmids were transformed into BL21 (DE3) Star *E. coli* strain. Bacterial cultures were grown in of LB/kanamycin (25 µg/mL) media at 37°C until reaching an OD of 0.6 (at a 600 nm wavelength). Protein expression was induced with 1 mM isopropyl 1-thio-β-D-galactopyranoside (IPTG) at 18 °C overnight. Cells were sonicated in ice and treated with benzonase (Merck-Millipore). Proteins were purified using immobilized metal ion affinity chromatography (IMAC) in a HiTrap TALON column (GE-Healthcare Life Sciences) with two washing steps: 50 mM buffer sodium phosphate (pH 7), 300 mM NaCl, and 50 mM buffer sodium phosphate (pH 7), 800 mM NaCl, before a 10-150 mM imidazole elution gradient. Removal of the histidine-tag was performed with GST-tagged PreScission Protease (GE-Healthcare Life Sciences). Further purification was performed using a HiTrap TALON column and a GST TALON column (GE-Healthcare Life Sciences). Purity and homogeneity were checked by SDS-PAGE and size-exclusion chromatography. Proteins were stored in buffer 50 mM Tris (pH 7.0) at −80 °C. The identity of all proteins was checked by mass spectrometry (4800plus MALDI-TOF/MS, from Applied Biosystems - Thermo Fisher Scientific). Stability and binding assays were performed at different pH and buffer conditions [50 mM Tris (pH 7-9), 0-150 mM NaCl; 50 mM Pipes (pH 7); 50 mM Phosphate (pH 7)]. Buffer exchanges were performed using a 10 kDa-pore size ultrafiltration device (Amicon centrifugal filter, Merck-Millipore) at 4000 rpm and 4 °C.

### Double-stranded DNA

HPLC-purified single-stranded DNA fragments were purchased from Integrated DNA Technologies. Sequences correspond to the promoter IV of the mouse brain-derived neurotrophic factor (BDNF) (16): forward unmethylated: 5’ -GCCATGCCCTGGAACGGAACTCTCCTAATAAAAGATGTATCATTT-3’; reverse unmethylated: 5’ -AAATGATACATCTTTTATTAGGAGAGTTCCGTTCC-AGGGCATGGC-3’; forward mCpG: 5’ -GCCATGCCCTGGAA(5-Me)CGGAACTCTCCTAATAAA-AGATGTATCATTT-3’; reverse mCpG: 5’ -AAATGATACATCTTTTATTAGGAGAGTTC(5-Me)CGTTCCAGGGCATGGC-3’. They were annealed using a Stratagene Mx3005 P qPCR real-time thermal cycler (Agilent Technologies) using the following thermal program: 1) 25 °C for 30 s; 2) 99 °C for 60 s; 3) cooling process down to 25 °C at a rate of 1 °C/180 s.

### Fluorescence spectroscopy

Thermal unfolding studies were performed in a Cary Eclipse fluorescence spectrophotometer (Varian – Agilent) in three steps using a 1 cm path-length quartz cuvette (Hellma Analytics). Temperature was controlled by a Peltier unit and monitored using a temperature probe. Fluorescence emission spectra were recorded from 300 to 400 nm using an excitation wavelength of 290 nm and a bandwidth of 5 nm. Protein concentration was set at 5 µM. Thermal stability assays were performed at a heating rate of 1 °C/min and at the wavelength for maximal spectral change (330 nm). Thermal unfolding experiments were analyzed considering a two-state unfolding model (16). The stabilizing effect upon dsDNA interaction was assessed performing thermal denaturations of the different proteins (at 5 µM) in the presence of methylated and unmethylated DNA (at 10 µM) under the same conditions.

### Isothermal titration calorimetry (ITC)

Proteins-dsDNA interactions were characterized using an Auto-iTC200 microcalorimeter (MicroCal – Malvern Instruments). Protein in the calorimetric cell at 3-5 µM was titrated with 50 µM dsDNA. All solutions were degassed at 15 °C for 2 min before each assay. A sequence of 2 µL-injections of titrant solution every 150 s was programmed and the stirring speed was set to 750 rpm. Interaction parameters (dissociation constant and binding enthalpy) were obtained as previously described (16). Buffer-independent binding parameters (*K*_*d*_, Δ*G*, Δ*H*, -*T*Δ*S*) and the number of protons released upon complex formation (*n*_*H*_) were determined by performing calorimetric titrations using buffers with different ionization enthalpies (16), and the binding heat capacity was determined by performing calorimetric titrations at different temperatures.

### Cell lines

Human neuroblastoma cell line SH-SY5Y was obtained from the American Type Culture Collection (ATCC No. CRL-2266) and were maintained in DMEM supplemented with 10% FBS, penicillin-streptomycin and L-glutamine (Gibco-BRL) in a 37°C, 5% CO_2_ humidified incubator. Differentiation was performed using a sequential treatment of Retinoic Acid (Sigma-Aldrich) and BDNF (Alomone Labs) as previously described (19). For transfection experiments cells were seeded and one day later were transfected with MeCP2-E1 and MeCP2-E2 C-terminal GFP fusion expression vectors (6) using Lipofectamine 3000 (Invitrogen) according to manufacturer instructions. Cycloheximide treatments were performed 48 h after transfection.

Cortical primary neurons were prepared from Sprague-Dawley rat embryos at gestational day 17 as described previously (72). If needed, they were transfected using the Amaxa rat neuron nucleofector kit (Lonza) following the manufacturer instructions (5.10^6^ neurons with 3 µg of plasmid DNA, by using the G-13 program on nucleofector apparatus) by plasmids expressing Flag-MeCP2 isoforms, kind gift from Angus Wilson (73). They were maintained in Neurobasal medium (Invitrogen) supplemented with 100 µg/ml penicillin/streptomycin, 2 mM glutamine, 2 % B-27 supplement (Invitrogen) during 7 to 9 DIV before proceeding to the depolarization or CHX assay. For depolarization assay, neuronal activity was blocked by pre-treatment of the cells with 1 mM tetrodotoxin (TTX), 100 mM DL-2-amino-5-phosphonovalerate (APV) and 20 mM 6-cyano-7-nitroquinoxaline-2, 3-dione (CNQX). Neuronal depolarization was subsequently achieved by 30 minutes exposure to 55 mM KCl in Tyrode buffer then neurons were incubated back in their culture medium during various times (17) before lysis.

### Cycloheximide chase assay

Cells were treated with 10 µg/mL of cycloheximide (Sigma-Aldrich) and harvested at the indicated times. Samples collected at each time point were then analyzed by western blot.

### Mass spectrometry to determine MeCP2 PTMs

All protein samples were digested overnight at 37°C with trypsin, using 50:1 protein:enzyme ratio. Digested peptide mixtures were desalted using C_18_ reverse phase columns, and then loaded onto a 50 cm x 75 μm ID column with RSLC 2 μm C_18_ packing material (EASY-Spray, Thermo-Fisher) with an integrated emitter, and then eluted into a Q-Exactive™ Hybrid Quadrupole-Orbitrap™ mass spectrometer (Thermo-Fisher) using an Easy-Spray nLC 1000 chromatography system (Thermo-Fisher,). The mass spectrometer was operated with 1 mass spectrometry (MS) spectrum followed by 10 MS/MS spectra in a data-dependent mode. The MS was acquired with a resolution of 70,000 FWHM (full width at half maximum), a target of 1 x 10^6^ ions and a maximum scan time of 120 ms. Using a relative collision energy of 27%, the MS/MS scans were acquired with a resolution of 17,500 FWHM, a target of 1 x 10^6^ ions and a maximum scan time of 120 ms. A dynamic exclusion time of 15 seconds was used for the MS/MS scans. XCalibur 2.2 (Thermo-Fisher Scientific) was used to acquire the raw data files and further processed with the PEAKS 7 search engine (Bioinformatics Solutions, Waterloo, ON) using a database consisting of the wild type MeCP2 constructs. Ascores were assigned for the peptides and PTMs using the PEAKS 7 software.

### Chromatin immunoprecipitation

Two months old mice were euthanized by exposure to CO_2_ 6 and 18 hours after first light stimulus (corresponding to 12 a.m. and 12 p.m. respectively). Brains were rapidly removed, dissected snap frozen and kept at −80°C until further use. One frontal cortex was grinded in 1 x PBS supplemented with complete protease inhibitor cocktail (Roche) and crosslinked in 1x PBS 0.5% formaldehyde at room temperature for 5 min. The crosslinking reaction was stopped by incubating the samples in 0.125 M Glycine for 5 min. The following steps were performed at 4 ºC. The tissue was centrifuge at 1500 x g for 5 min and washed twice with ice cold PBS using similar centrifugation steps. The pellet was resuspended in 1 mL ChIP lysis buffer [10 mM HEPES (pH7.9), 1.5 M MgCl_2_, 10 mM KCl, 0.5 mM DTT, 0.1% NP-40] supplemented with cOmplete Protease Inhibitor Cocktail (Roche), dounce-homogenized (10 strokes) and incubated on ice for 20 min. Nuclei were centrifuged at 2500 x g for 5 min and resuspended in 300 µL of RIPA buffer [50 mM Tris-HCl (pH 8), 150 mM NaCl, 0.5 % SDS, 0.5 % Sodium deoxycholate, 1 % Triton-100] supplemented with cOmplete Protease Inhibitor Cocktail (Roche). Suspension was then sonicated in a Bioruptor (Diagenode) at high power for 15 min with 30 seconds on/off intervals. Chromatin was centrifuged at 16000 x g for 10 min. Supernatant was diluted 10 times with RIPA buffer without SDS [50 mM Tris-HCl (pH8.0), 150 mM NaCl, 0.5% Sodium deoxycolate, 1% Triton-100] and pre-cleared for 2 hours with 20 µL of Dynabeads Protein G magnetic beads (Thermo Fisher Scientific). Eight µg of MeCP2 E1 and E2 antibodies (produced in house, Supplementary Fig. 4A) and Normal Rabbit IgG (Cell Signaling Technologies) were bound to 25 µL of Dynabeads Protein G following the manufacturer instructions and incubated 1 hour in PBS/5% BSA. Five per cent of this chromatin was kept aside as Input sample. The remaining chromatin was aliquoted in 3 tubes and incubated with the corresponding antibody at 4 ºC overnight while tumbling. Next day the supernatants were discarded and the beads were washed twice with 1 mL of low salt buffer [50 mM Tris-HCl (pH 8.0), 150 mM NaCl, 0.1% SDS, 1% NP-40, 1 mM EDTA, 0.5% Sodium deoxycolate], twice with 1 mL of high salt buffer [50 mM Tris-HCl (pH 8.0), 500 mM NaCl, 0.1% SDS, 1% NP-40, 1 mM EDTA, 0.5% Sodium deoxycolate], twice with 1 mL of LiCl buffer [50 mM Tris-HCl (pH 8.0), 250 mM LiCl, 0.1% SDS, 1% NP-40, 1 mM EDTA, 0.5% Sodium deoxycolate] and twice with 1 mL of TE. Chromatin was eluted twice by addition of 100 µL of elution buffer (100 mM NaHCO_3_, 1% SDS) and 10 min incubations at 65 ºC. The volume of the input sample was adjusted to 200 µL with elution buffer and reversal of the crosslinking was carried out for 5 hours at 65 ºC. The samples were then incubated with proteinase K for 1 hour at 65 ºC and RNase A for 30 min at 42 ºC. DNA was purified with a PCR purification kit (Qiagen).

### Library preparation and sequencing

All liquid handling steps were carried out on the Bravo liquid handling platform using VWorks Automation Control Software (Agilent Automation). Enzyme mastermix dispenses and magnetic bead clean up steps were performed using the Bravo configured with the 96LT pipetting head (Agilent Technologies). End Repair and 5’ Phosphorylation reaction [35 µL DNA sample, 5 µL of 10X NEB 2 buffer, 2 µL of 25 mM ATP, 2 µL of 10 mM dNTP, 10U T4 Polynucleotide Kinase, 4.5U T4 DNA Polymerase, 1U Klenow Large Fragment DNA Polymerase, and ultrapure water (New England Biolabs)] were incubated for 30 minutes at room temperature. DNA samples were then purified using house-made magnetic bead solution [1M NaCl, 23% PEG, Sera-Mag Speedbeads (Thermo Fisher Scientific)] with final PEG concentration of 13.87%. DNA was eluted with Qiagen EB buffer. A single dA overhang was added to the 3’ ends of DNA fragments [35 µL DNA, 5 µL of 10X NEB 2 buffer, 1 µL of 10 mM dATP, 5U Klenow Fragment (3′ →5′ exo–), and ultrapure water (New England Biolabs)]. The product was then purified using house-made magnetic bead solution [1M NaCl, 23% PEG, Sera-Mag Speedbeads (Thermo Fisher Scientific)] with final PEG concentration of 13.87%. Next, Illumina short sequencing adaptors were ligated to the dA-tailed DNA fragments. Ligation products were then purified two times using house-made magnetic bead solution with final PEG concentrations of 8.89% and 10.91% respectively. Adaptor ligated libraries were PCR amplified and barcoded by custom indexing primers in a 60 µL PCR reaction [35 µL DNA, 12 µL of 5X High fidelity buffer, 1 µL of 10mM dNTPs, 1.5 μL of DMSO, 1 µL of 25 μM PCR primer 1.0, 2 µL of 12.5 µM custom indexing primer, 1U Phusion Hot Start II, and ultrapure water to a total volume of 60 µL (Thermo Fisher Scientific)]. PCR amplified libraries were size selected with house-made magnetic bead solution [1M NaCl, 20% PEG, Sera-Mag Speedbeads (Thermo Fisher Scientific)] with final PEG concentration of 9.19%. The average size was determined by running 1 µL of each final library product on the Agilent HS DNA assay chip. Libraries were quantified using Qubit HS DNA assay with Qubit Fluorometer (Thermo Fisher Scientific). Library pool was sequenced paired-end 75 on HiSeq 2500 (Illumina) and aligned to a mouse genome (mm10) using BWA 0.5.7.

### Bioinformatics analysis of ChIP-Seq data

The reads were first trimmed with sickle (74) to remove low quality ends, then mapped with BWA (75) on the GRCm38 mouse assembly. Reads mapping exactly once as proper pairs were kept. Since the signal was unclear on each individual replicate, we decided to merge them. We detected binding sites using SICER (window 600 bp, gap 200 bp) (32), and we also detected differential binding sites (*e.g.* E1 vs E2) with the same tool. RSAT peak-motif tool (76) was used detect over-represented motifs in the detected peaks, and ChIPpeakAnno (77) was used to annotate the peaks. With deepTools (33), we first normalized the samples in order to have similar RPKM per bin, and computed the fold change IP over input. We computed the fold change distribution along the gene body, the TSS and the TES, using GRCm38.0 as reference annotation. We then computed the fold change distribution along each gene, and clustered the genes into 5 groups using the k-means method, where genes in each group tend to have the same fold change distribution. Pathway enrichment analysis was performed with WebGestalt (78).

### RT-qPCR

Each PCR was carried out in triplicate using SYBR Green PCR Master Mix (Applied Biosystems). Thermocycling conditions were 10 minutes at 95°C, then 40 cycles of 15 seconds at 95°C and 1 minute at 60°C. Fluorescent signals were acquired by the Stratagene Mx3005P qPCR System (Agilent Technologies). Primer sequences are as follows: Fw Olfr52: GCCCAGTTAGGCTGCTTTCT and Rv Olfr52: CAAGCTGAATGCAGATTCCA; Fw Olfr786: ACTCCCTTGGTCAATGATGC and Rv Olfr786: GCCTTTTCCACATGCTCTTC; Fw Olfr1510: ACCTGCTCATCCTGCTCACT and Rv Olfr1510: AGGAGAGCACACCCAGAAGA; Fw Rps3: GCAGCGTAGAGGTGAGTTCC and Rv Rps3: ACTGGCCATGTCAGGTTTTC; Fw Hist1h4b: AGGAACACCTTCAGCACACC and Rv Hist1h4b: GACAACATCCAGGGCATCAC; Fw Rpl7a: GGACTTTGAGCCGCTTGTAG and Rv Rpl7a: GAGATTTAACGCGCTTCGTC.

### Co-immunoprecipitation and mass spectrometry

One whole brain was dounce-homogenized in buffer A [20 mM HEPES (pH7.5), 10 mM KCl, 1.5 mM MgCl_2_, 0.34 M Sucrose, 10 % glycerol, 1 mM DTT, 0.5 % Triton] [All buffers supplemented with cOmplete Protease Inhibitor Cocktail EDTA-free and PhosSTOP (Roche) and all steps performed in ice unless otherwise indicated]. Tissue was incubated 10 min and centrifuged. Supernatant was discarded and nuclei pellet washed with MNase digestion buffer [15 mM NaCl and 50 mM Tris-HCl (pH 7.5)]. Pellet was resuspended in digestion buffer with 2 mM CaCl_2_ and incubated for 30 min at 37ºC with 150 ud MNase (Worthington). After centrifugation, supernatant was kept on ice and reaction was stopped by addition of EDTA (10 mM final concentration). The remaining pellet was resuspended in digestion buffer with 2 mM CaCl_2_, 50 ud MNase were added and incubated for 20 min at 37ºC. The reaction was stopped by addition of EDTA (10 mM final concentration), the sample was centrifuged and supernatant kept. The remaining pellet was resuspended in TE buffer [10 mM Tris-HCl (pH 8.0), 0.25 mM EDTA] and kept tumbling for 30 min. All fractions were combined, centrifuged at maximum speed during 10 minutes and the pellet was discarded. Salt adjustment of the lysate was performed by addition of 3x buffer D [60 mM HEPES (pH 7.5), 450 mM NaCl, 4.5 mM MgCl_2_, 0.6 mM EGTA, 0.6 % Triton, 30 % glycerol] dropwise with gentle vortexing. Lysate was centrifuged 10 min at maximum speed and supernatant was precleared for 1 hour with 50 µLof Dynabeads Protein G magnetic beads (Thermo Fisher Scientific). Negative controls for the IP were obtained by blocking E1 and E2 paratopes with E1 and E2 specific peptides [E1: CAAAAPSGGGGGGEEER; E2: MVAGMLGLREEKC (New England Peptides)]. Antibodies were blocked during 45 min by tumbling at room temperature with a 5 fold excess of the specific peptide. 10 µg of each antibody [E1, E1 blocked, E2, E2 blocked and normal rabbit IgG (Cell Signaling Technologies)] were bound to 50 µl of magnetic beads according to manufacturer instructions. Immunoprecipitations were carried out overnight at 4ºC while tumbling. Antibody-protein complexes were washed 3 times with 1x buffer D supplemented with 0.5 % Triton. Proteins were eluted with 2x SDS buffer, boiled for 10 min and proteins were run on SDS-PAGE. The gel was stained with Coomassie blue and different sections were excised for subsequent analysis. Mass spectrometric analysis was then performed using a nano-HPLC system (Easy-nLC II, Thermo Fisher Scientific), coupled to the ESI-source of an LTQ Orbitrap Velos (Thermo Fisher Scientific), using conditions described previously (79). Briefly, samples were injected onto a 100 μm ID, 360 μm OD trap column packed with Magic C18AQ (Bruker-Michrom), 100 Å, 5 μm pore size (prepared in-house) and desalted by washing with solvent A [2% acetonitrile:98% water, 0.1% formic acid (FA)]. Peptides were separated with a 60-min gradient [0-60 min: 5-40% solvent B (90% acetonitrile, 10% water, 0.1% FA), 60-62 min: 40-80% B, 62-66 min: 80-100% B, 66-70 min: 100% B], on a 75 μm ID, 360 μm OD analytical column packed with Magic C18AQ 100 Å, 5 μm pore size (prepared in-house), with IntegraFrit (New Objective Inc.) and equilibrated with solvent A. MS data were acquired using a data dependant method. The data dependent acquisition also utilized dynamic exclusion, with an exclusion window of 10 ppm and exclusion duration of 10 seconds. MS events used 60000-resolution FTMS scans, and MS/MS events used ITMS scans, with a scan range of m/z 400-2000 in the MS scan. MS/MS data were analyzed using Mascot. The data was compared to the Uniprot Mouse database, using trypsin digestion with up to 3 missed cleavages, a peptide tolerance of 5 ppm, and MS/MS tolerance of 0.3 Da. Acetylated N-termini and oxidation of methionine were included as variable modifications.

## Acknowledgments

This work was supported by grants from the Canadian Institutes of Health Research; (CIHR grant MOP-130417) to JA. Genome Canada and Genome British Columbia for financial support for the University of Victoria-Genome BC Proteomics Centre (project codes 204PRO for operations and 214PRO for technology development), Leading Edge Endowment Fund (University of Victoria), the Segal McGill Chair in Molecular Oncology at McGill University (Montreal, Quebec, Canada), the Warren Y. Soper Charitable Trust and the Alvin Segal Family Foundation to the Jewish General Hospital (Montreal, Quebec, Canada) CHB. Max Planck Society and DFG CRC 1080; DFG CRC 902 and European Research Council (ERC) under the European Union’s Horizon 2020 research and innovation programme (grant agreement No 743216) to EMS. CIHR grant MOP-102758 to JBV. Spanish Ministerio de Economia y Competitividad [BFU2013-47064-P and BFU2016-78232-P] to AVC. Instituto de Salud Carlos III and co-funded by European Union (ERDF/ESF, “Investing in your future”) [PI15/00663, and Miguel Servet contract CPII13/0017] to OA. Spanish Ministerio de Educacion Cultura y Deporte [FPU 13/3870] to RCG. Terry Fox Research Institute Program Project (Grant: TFRI #1039) and Canadian Cancer Society Research Institute (Grant: CCSRI #703489) to MH. European Research Council under the European Union’s Horizon 2020 research and innovation programme under grant agreement No 727264 (EPIPHARM); the European Community’s Seventh Framework Programme (FP7/2007–2013)/ERC Grant Agreement 268626/EPINORC project; the Ministerio de Economía y Competitividad (MINECO) under grant no. SAF2014-55000-R; the Integrated Project of Excellence no. PIE13/00022 (ONCOPROFILE); the CIBER 2016 CB16/12/00312 (CIBERONC); the Spanish Cancer Research Network (RTICC) no. RD12/0036/0039; the Cellex Foundation; Obra Social “La Caixa”; and the Health and Science Departments of the Catalan Government (Generalitat de Catalunya) AGAUR– project no. 2014SGR633 to ME. ME is an ICREA Research Professor. We are also thankful to the French Association of Rett Syndrome (AFSR) that funded research conducted by CEM and HM and funded LK, genotoul bioinformatics platform Toulouse Midi-Pyrenees (Bioinfo Genotoul) for providing computing and storage resources.

AUTHOR CONTRIBUTIONS
AMdP and JA designed and wrote the paper. AMdP, MSC, HM, CM, MEF and TIS performed the biochemical/molecular biology experiments, RC-G, OA and AV-C designed and performed the calorimetry and spectroscopy experiments, MMM provided assistance with construction of the ChIP libraries, NIB and EVP carried out the proteomic analyses, LK, MZ and AC conducted the bioinformatic analyses of the CHIP-seq data, STD and EMS performed the immunofluorescence, JVS-M, MZ, AV-C, JBV, MH, ME and CM contributed to the writing and discussion of specific sections of the manuscript.

